# Multi-omics approach highlights differences between functional RLP classes in *Arabidopsis thaliana*

**DOI:** 10.1101/2020.08.07.240911

**Authors:** C. Steidele, R. Stam

## Abstract

The receptor-like protein (RLP) family is a complex gene family with 57 members in *Arabidopsis thaliana*. Some members of the RLP family are known to be involved in basal developmental processes, whereas others have found to be involved in defence responses. However, functional data is to date, only available for a small subset of RLPs, leaving the remaining ones classified as RLPs of unknown function. Using publicly available datasets, we annotated those RLPs of unknown functions as either likely defence-related or likely fulfilling a more basal function in plants. Using these categories, we can identify important characteristics that differ between the RLP sub classes. We find the two classes differ in abundance on both transcriptome and proteome level, physical clustering in the genome and putative interaction partners. However, the classes do not differ in the genetic diversity of their individual members in accessible pan-genome data. Our work has several implications for work related to functional studies on RLPs as well as for the understanding of RLP gene family evolution. Using our annotations, we can make suggestions of which RLPs can be identified as potential immune receptors using genetics tools, which can be useful for disease studies. The lack of differences in nucleotide diversity between the two RLP subclasses further suggests that non-synonymous diversity of gene sequences alone cannot distinguish defence from developmental genes. By contrast, differences in transcript and protein abundance or clustering at genomic loci might also allow for functional annotations and characterisation in other plant species.

## Introduction

Plants are caught in ever ongoing evolutionary interactions with their pathogens, that have, dependent on their nature, been described as arms races or trench warfare, each with their own underlying evolutionary dynamics (Tellier *et al*. 2014). In either case plants need to evolve resistance mechanisms in order to survive, at the same time, pathogens need to evolve to overcome these resistances to be able to infect, which leads to the necessity of the plant’s defences to evolve again. This leads to the hypothesis that defence associated genes should be faster evolving than, for example, development associated genes. On a phylogenetic scale this can be illustrated by very large, diverse and expanded resistance associated gene-families. Most known are the intracellular receptor genes of the NLR family (nucleotide-binding domain and leucine-rich repeat containing). This family, but also other Leucine-rich repeat (LRR) containing defence-associated genes, drastically diversified over the course of evolution. Indeed NLRs are much more diverse than for example the defensin gene family, that is known to have dual roles in defence as well as development (Mondragón-Palomino *et al.* 2017). The enormous variation in NLRs between species and also variation in how these modular receptors are built-up have been discussed in many different papers (Monteiro and Nishimura 2018; Stam *et al.* 2019a).

How much diversity exists in defence gene families within a species is a lesser studied topic. Recently polymorphisms and significant copy number variations have been documented within the NLR family in 64 resequenced *Arabidopsis thaliana* accessions (Van de Weyer *et al*. 2019) and a study investigating sequence polymorphisms in NLRs in a single tomato species found that NLRs experience different selective pressures dependent on the geographical location of the population (Stam *et al.* 2019b). These studies thus highlight that indeed defence associated gene families appear to be highly diverse, but do not allow comparisons between defence and development associated genes in the same gene family. Besides the NLRs, plants have evolved a plethora of plasma-membrane bound or associated receptors to monitor their environment, but also as a communication tool within the plant itself to facilitate for example stomatal patterning. The different plasma-membrane located receptors can be divided into two major groups, receptor-like kinases (RLKs) with an intracellular signalling domain and receptor-like proteins (RLPs), which only contain a small or no cytoplasmic tail. Besides the differentiation between RLKs and RLPs, the receptors can be categorized according to their extracellular domains. These domains can facilitate binding and recognition of the corresponding ligands or enable interaction with other proteins to maintain or finetune signalling. In Arabidopsis more than 600 RLKs are annotated (Shiu and Bleecker 2001) and 57 LRR-RLPs are identified and numbered in consecutive order according to their gene numbers along the Arabidopsis genome (Wang *et al*. 2008a; Fritz-Laylin *et al.* 2005). Members of the LRR-RLP family have been shown to be involved in both developmental and defence mechanisms, making them ideally suited to investigate whether functional differences lead to differences in rates of evolution.

Of the 57 annotated RLPs in Arabidopsis thaliana 2 RLPs are experimentally validated to be associated with developmental functions (RLP10/CLV2, RLP17/TMM), and 6 with defence functions (RLP1, 3, 23, 30, 32, 42). CLAVATA2(CLV2)/RLP10 seems to be a unique RLP as it plays a role both in developmental and defence-related processes. The best characterised function of CLV2 is the regulation of the shoot apical meristem (SAM) maintenance, but it also plays a role in the regulation of the root apical meristem (RAM) maintenance, the regulation of the protoxylem formation, organ development and plant-microbe interactions (Pan *et al*. 2016). Additionally, two other RLPs (RLP2 and 12) can rescue the *clv2*-phenotype, when the corresponding genes are expressed under the clv2-promoter (Wang *et al*. 2010). RLP17, also named TOO MANY MOUTH (TMM), is involved in the regulation of the patterning of stomata, micropores to facilitate gas exchange which are located in the epidermis of plant leaves (Nadeau and Sack 2002; Shpak *et al.* 2005).

Fritz-Laylin *et al*. (2005) used a comparative approach with several criteria, like global alignability, genomic organization and sequence identity to identify PUTATIVE DEVELOPMENTAL ORTHOLOGS (PDOs) in Arabidopsis and rice. Based on this classification 4 RLPs could be identified: PDO1/RLP51, PDO2/RLP4, PDO3/RLP10/CLV2, PDO4/RLP17/TMM. Furthermore, they could show that based on phylogenetic comparisons 47 of 57 AtRLPs group together in so-called superclades. They found that the PDOs do not fall into those superclades, nor do RLP29, 44, 46, 55, 57. Thus for these RLPs a putative function in development was hypothesized (Fritz-Laylin *et al.* 2005). It was later shown that RLP44 is mediating the response to pectin modification by activating brassinosteroid signaling (Wolf *et al.* 2014) and is important for the regulation of xylem fate (Holzwart *et al*. 2018). PDO1/RLP51 is the underlying gene of the snc2-1D locus (for suppressor of npr1, constitutive 2-1D), a semidominant gain-of-function *Arabidopsis thaliana* mutant with constitutively activated defense responses, dwarf morphology and high salicylic acid and PATHOGENESIS-RELATED (PR) genes levels (Zhang *et al.* 2010). Therefore, we refer to those 9 RLPs (RLP4, 10/CLV2, 17/TMM, 29, 44, 46, 51, 55, 57) as PDOs.

Several RLPs have been shown to fulfill important roles in the defence against pathogens. Plants have evolved a two-layered, pathogen-activated immune system to detect and fight off invading pathogens: pattern-triggered immunity (PTI) or surface immunity and effector-triggered immunity (ETI) or intracellular immunity. According to the current and simplified paradigm, pathogen associated molecular patterns (PAMPs) are recognized by cell-surface localized pattern recognition receptors and larger pathogen-secreted proteins, called effectors, are typically recognized by intracellular NLR-receptors (Jones and Dangl 2006; Boller and Felix 2009; Dodds and Rathjen 2010; van der Burgh and Joosten 2019). There is some debate as to whether the separation of the recognised molecules (PAMPS vs effectors) can be made that strictly (Thomma *et al.* 2011; van der Burgh and Joosten 2019), several LRR-RLPs have been validated to be able to facilitate immune responses to help protecting the plant against different pathogens.

RLP1/ReMAX (RECEPTOR of eMAX) can detect the ENIGMATIC MAMP OF XANTHOMONAS (eMAX) (Jehle *et al*. 2013a, 2013b), RLP23 detects a widespread, but conserved twenty amino acid long epitope in NECROSIS AND ETHYLENE INDUCING (NEP) - LIKE PROTEINS (NLPs) (Albert *et al.* 2015). This so-called nlp20 motif is present in NLPs from fungi, oomycetes and bacteria (Böhm *et al*. 2014b). A still unidentified SCLEROTINIA CULTURE FILTRATE ELICITOR 1 (SCFE1) is perceived via RLP30 (Zhang *et al*. 2013), and the same RLP additionally detects a bacterial elicitor called PSEUDOMONAS CULTURE FILTRATE ELICITOR 1 (PCFE1) (Feiler 2020). RLP32 recognizes the structural fold of the bacterial translation initiation factor -1 (Inf-1) present in all proteobacteria (Melzer 2013; Fan 2016) and RLP42/RBPG1 detects several endopolygalacturonases from *Botrytis cinerea* and *Aspergillus niger* (Zhang *et al*. 2014). Finally, RLP3 is the causal gene of the quantitative resistance locus RFO2 in Arabidopsis conferring resistance against the vascular wilt fungus *Fusarium oxysporum* forma specialis *matthioli* (Shen and Diener 2013). As these 6 RLPs (RLP1, 3, 23, 30, 32 and 42) are validated to play important roles in the defence against various pathogens we will refer to them as VDRs (validated defence RLPs) in the remainder of this manuscript.

RLPs lack an obvious intracellular signalling domain and thus require additional interaction partners. For the VDR RLP23 it was shown that the short cytoplasmic tail has if only an auxillary, but not essential function in nlp20-mediated ethylene signalling *(Albert et al.* 2019). The VDRs RLP1, RLP23, RLP30, RLP32 and RLP42 all require BRASSINOSTEROID-INSENSITIVE KINASE 1 (BAK1) and SUPPRESSOR OF BIR1 (SOBIR1) for full function.The mentioned RLPs are constitutively interacting with SOBIR1 at the plant plasma membrane and upon ligand perception BAK1 is recruited to the complex (Jehle *et al*. 2013b; Albert *et al*. 2015; Zhang *et al*. 2013; Fan 2016; Zhang *et al*. 2014).The interaction with SOBIR1 is mediated via a stretch of negatively charged amino acids, Aspartate (D) and Glutamate (E), in the extracellular juxtamembrane region, just before the transmembrane domain and a conserved GxxxG motif within the transmembrane region (Albert *et al.* 2019).

The PDO RLP10/CLV2 interacts with the kinase CORYNE (CRN) and together they can form a multimer with the LRR-kinase CLAVATA 1(CLV1) (Somssich *et al*. 2016). And RLP17/TMM forms a receptor complex with the ERECTA RECEPTOR KINASES (ER) or ER-LIKE 1 (ERL1) to regulate stomatal patterning (Lin *et al.* 2017). Whereas these analyses are far from complete, they seem to suggest distinct evolutionary trajectories for PDOs and VDRs.

Over the last decades, a large number of publicly available datasets have become available for *A. thaliana* research. These data sets range from (reference) genome data (The Arabidopsis Information Resource, TAIR, Berardini *et al*. 2015) and gene expression atlasses (Hruz *et al.* 2008) to the 1001 Arabidopsis genome project (Alonso-Blanco *et al*. 2016). Very recently a full *A. thaliana* transcriptome and proteome database has been published (Mergner *et al.* 2020) as well as a copy number variant atlas, cataloging presence and absence variation between over 1100 *A. thaliana* accessions (Zmienko *et al.* 2020). The availability of these data sets for the first time allows to compare the diversity of genes and gene families on many levels.

In this paper we utilize these publically available datasets to gain a deeper understanding of the RLP family in Arabidopsis. Knowing that the RLP family contains both developmental and defence-associated members, we specifically focus on comparing those two classes. We investigate the two subfamilies on all levels, ranging from phylogenetic relationship to gene expression and species-wide polymorphisms. Our results show clearly distinguishable characteristics between defence and development associated RLPs.

## Results

### Phylogenetic clustering can infer RLP functions

First we wanted to know whether we could split the RLP family in a defence associated and a development associated fraction. The most straightforward way to infer RLP functions would be if genes with similar functions e.g. defence or conserved roles, would share higher sequence similarity and thus cluster together in phylogenetic trees. Four papers studied the phylogeny of the RLPs before (Tör *et al.* 2004; Fritz-Laylin *et al.* 2005; Mondragon-Palomino and Gaut 2005; Wang *et al.* 2008a), however, at the time of publication not many RLPs were functionally annotated. Figure 1 shows the most up to date phylogeny as constructed by Wang *et al.* 2008a using the conserved C3-F domain of the proteins. We used this tree to annotate the above-mentioned PDOs (RLP4, 10/CLV2, 17/TMM, 29, 44, 46, 51, 55, 57) and VDRs (RLP1, 3, 23, 30, 32 and 42). The PDOs, except RLP4 and RLP46 are all on one branches of the phylogenetic tree, whereas the defence associated RLPs are more scattered across the tree. This is in line with previous publications where already a higher number of RLPs was predicted to be associated with defence (Fritz-Laylin *et al.* 2005). It was shown that 47 out of the analyzed 57 RLPs cluster within superclades where at least one member was defence-associated.

**Figure 1:**
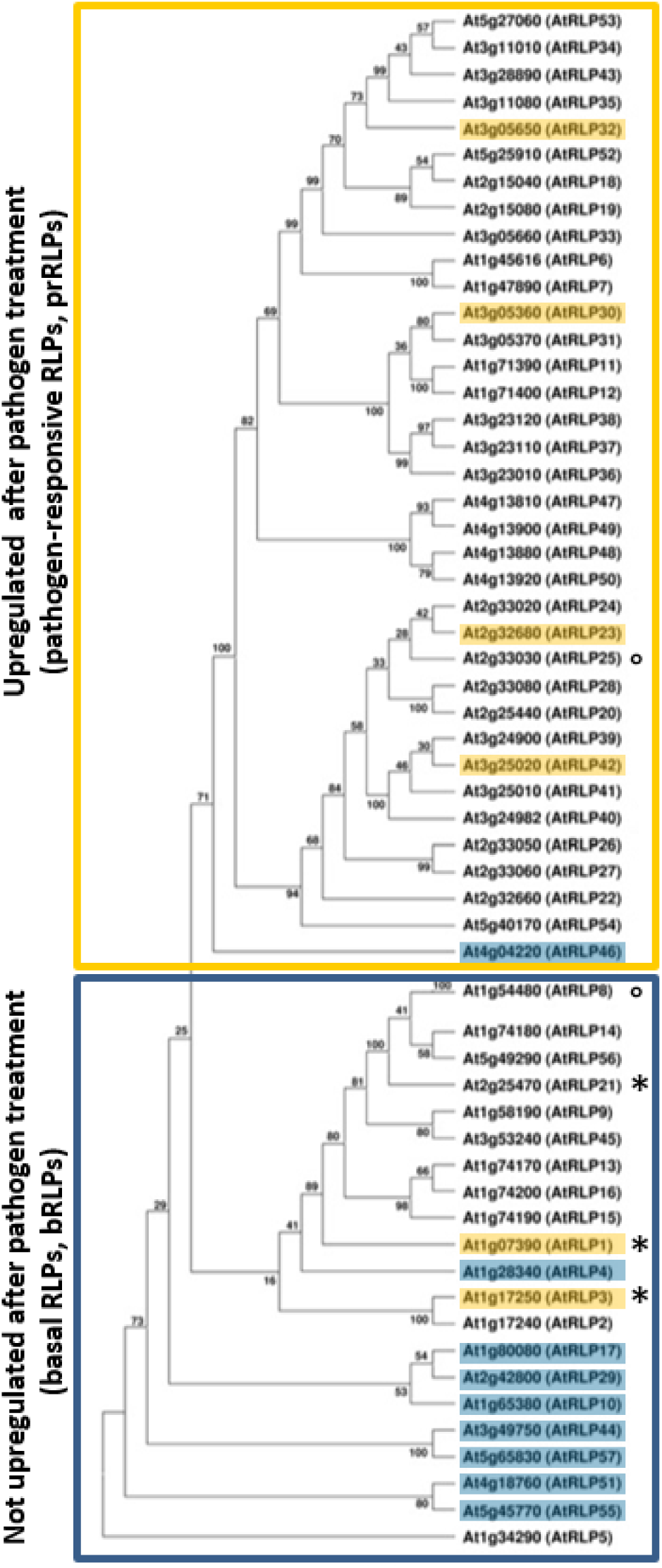
Phylogenetic tree of the conserved C3-F domain of RLPs, modified from Wang *et al.* 2008a. The bRLPs which are not upregulated after pathogen treatment and the prRLPs which are upregulated after pathogen treatment form distinct groups within the phylogenetic tree, resembling the already known or assumed functions of the PDOs and VDRs. Highlighted in blue are PDOs and in yellow VDRs. Boxed in yellow are the prRLPs that are at least 2.5x upregulated with a p-value of 0.001 after infection with various pathogens (except AtRLP6, 47 and 48 which are only 1.5x upregulated). Boxed in blue are the bRLPs which were not upregulated by pathogen infection (|2.5|, p-value=0.001). Used datasets are AT_AFFY_ATH1-0 and AT_mRNAseq_ARABI_GL-3. *AtRLP1, AtRLP3 and AtRLP21 showed an upregulation after pathogen treatment. °AtRLP25 is not up or down regulated at all and AtRLP8 was not present in the used datasets.

**Figure 2:**
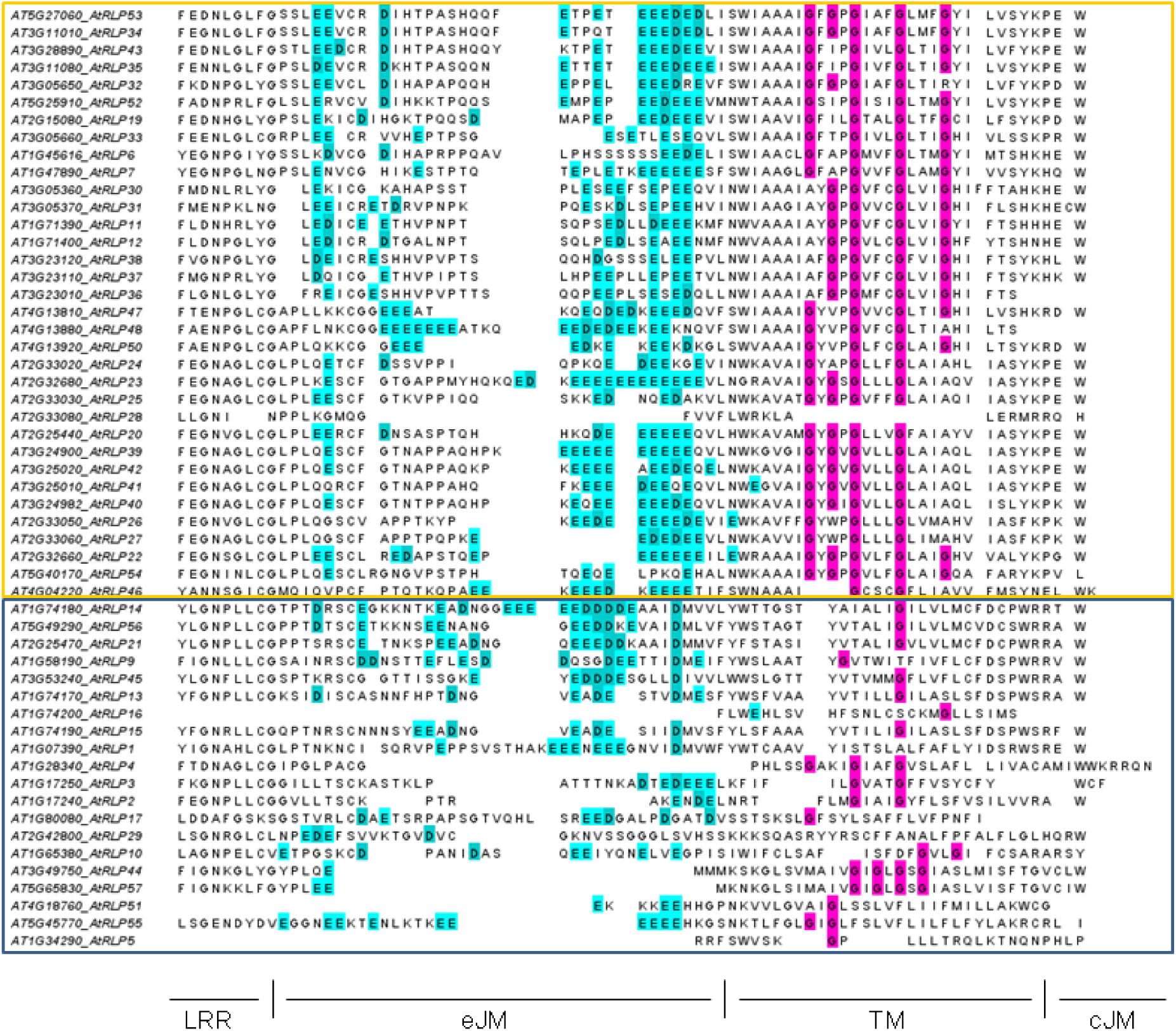
Alignment of the extracellular juxtamembrane, transmembrane and cytoplasmic region of the RLPs. Most of the prRLPs (boxed in yellow) have both motifs required for interaction with SOBIR1, the negatively charged amino acid stretch in the eJM and the GxxxG-motif in the TM, except RLP28 lacking both motifs and RLP46 having only two Gs. Most of the not-upregulated bRLPs (boxed in blue) lack the GxxxG-motif, except RLP44 and RLP57, but the latter lack a dominant negatively charged amino acid stretch in the eJM. All sequences were aligned using muscle and the sequences were ordered manually to fit the phylogenetic tree. The last Leucine rich repeat-domain (LRR), as well as the extracellular juxtamembrane (eJM), the transmembrane domain (TM) and the cellular juxtamembrane (cJM) are indicated. Color coded in magenta are the Glycines (G) in the TM and in cyan the Aspartates (D) and Glutamates (E) in the eJM.

### Expression data after induction confirmed different roles for basal RLPs and pathogen-responsive RLPs

Based on the findings above, we hypothesized that RLPs on the upper branches of the tree are likely to be defence associated. To expand the annotation data of the RLPs, we used the Genevestigator software (Hruz *et al*. 2008). The expression of those RLPs after pathogen treatment was checked in two different datasets containing expression data for treatment of *A. thaliana* with several bacterial and filamentous pathogens. 35 RLPs showed an upregulated gene expression after treatment with pathogens in at least one of the different pathogen infection datasets, whereas 17 RLPs showed no changes in expression after pathogen treatment in any of the examined data sets. As expected, all previously identified defence-associated RLPs are upregulated, whereas none of the PDOs show changes after infection. Interestingly, when we superimpose the expression data on the phylogeny we see a near perfect separation of upregulation in all higher branches and no regulation in the lower, basal branches (Figure 1). The only three exceptions are the two defence-associated RLPs mentioned before, RLP1 and RLP3, as well as RLP21. Only RLP25 shows no changes in expression in any of the examined datasets and RLP8 was missing from the data.

When combined, these data strongly suggest that the upper part of the phylogenetic tree completely consists of defence associated RLPs, that all derived from more ancestral basal, putative developmental-related RLPs. In the remainder of this manuscript we will therefore refer to the upper part of the phylogeny as prRLPs (pathogen-responsive RLPs) and the lower part as bRLPs (basal RLPs).

### prRLPs are species specific

Now that we have established that within Arabidopsis, RLPs can be clearly divided in two major groups, we wondered whether this division can also be seen outside the species. Kang and Yeom (2018) recently published a completely updated annotation of all RLPs in tomato (*Solanum lycopersicum)*. Similar to Wang *et al*. 2008a, they generated a phylogenetic tree from the C3-F domain, using all available RLPs for both tomato and Arabidopsis. Interestingly, the bRLPs can be found both as poly- and paraphyletic groups with the annotated tomato RLPs, the prRLPs all form a single monophyletic group (Figure S1), thus indicating that the prRLPs derive from species-specific family expansions.

**Figure S1:**
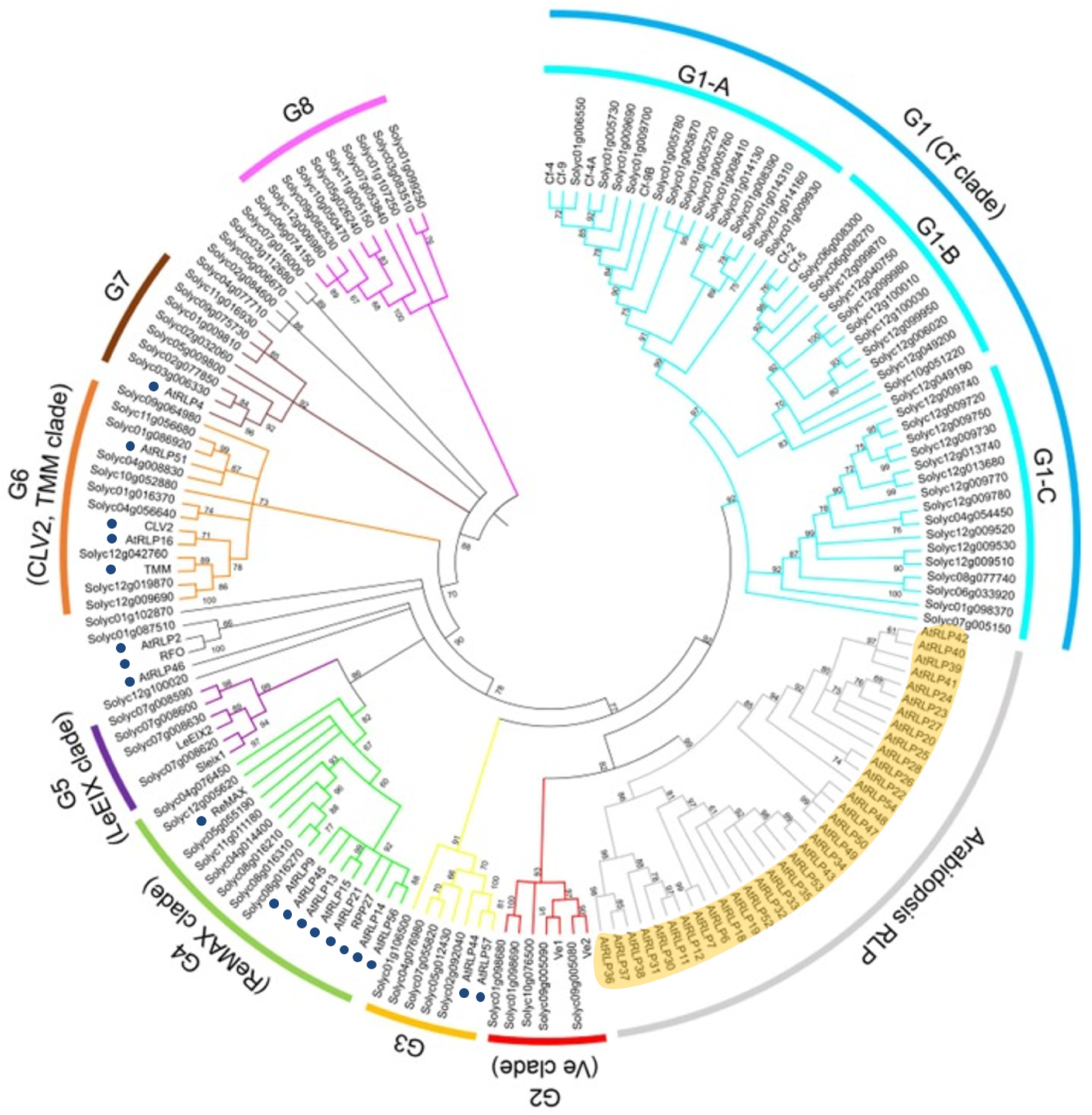
The phylogenetic tree of tomato and Arabidopsis RLPs, published by Kang and Yeom 2018, shows that the Arabidopsis prRLPs (highlighted in yellow) form a monophyletic clade, whereas the Arabidopsis bRLPs cluster as poly- and paraphyletic groups with the annotated tomato RLPs (marked with blue dots).

### bRLPs lack common protein-interacting motifs

Defence-associated RLPs constitutively interact with SOBIR1 (Bi *et al.* 2014; Albert *et al*. 2015) and for this interaction two motifs are important: a stretch of negatively charged amino acids in the extracellular juxtamembrane region and a conserved GxxxG-motif in the transmembrane domain (Albert *et al.* 2019).

All of the known prRLPs possess the conserved stretch of negatively charged amino acids (Aspartate D and Glutamate E). Only RLP1 lacks the GxxxG-motif, but it was shown that it can still interact with SOBIR1 (Albert *et al.* 2019). From the bRLPs only RLP46 and RLP55 have a pronounced stretch of Aspartate and Glutamate. RLP17/TMM and RLP29 contain neither the negatively charged amino acids nor the GxxxG-motif. We expanded these analyses and investigated the presence of these motifs in the complete prRLP set and the non regulated bRLP set and find that with only one exception all pathogen-responsive RLPs contain both motifs, whereas one or in some cases even both motifs appear to be absent in the bRLP set. This might suggest that SOBIR1-dependency evolved in relation to a function in pathogen defence.

### Basal and pathogen-responsive RLPs cluster in the transcriptome

Seeing that we can now assign defence responsive and basal functions to all RLPs, we wanted to know if besides phylogenetic separation the two groups show other globally different characteristics. For example, one can hypothesize that defence associated and non defence associated RLPs also show different transcript levels in non elicited states.

Therefore, we examined the steady state expression levels of all RLPs in different tissues. We obtained such expression data, consisting of different tissue samples from the Arabidopsis proteome project (Mergner *et al*. 2020) and looked for similar expression patterns using an hierarchical clustering method. Figure 3A shows clear clustering into a predominantly pathogen-responsive cluster (88% of the genes are prRLPs) and one cluster with mainly bRLPs (77% of the genes are bRLPs). It should be noted that no information was available for RLP5, RLP8, RLP11, RLP15, RLP18, RLP21, RLP25 and RLP49.

**Figure 3:**
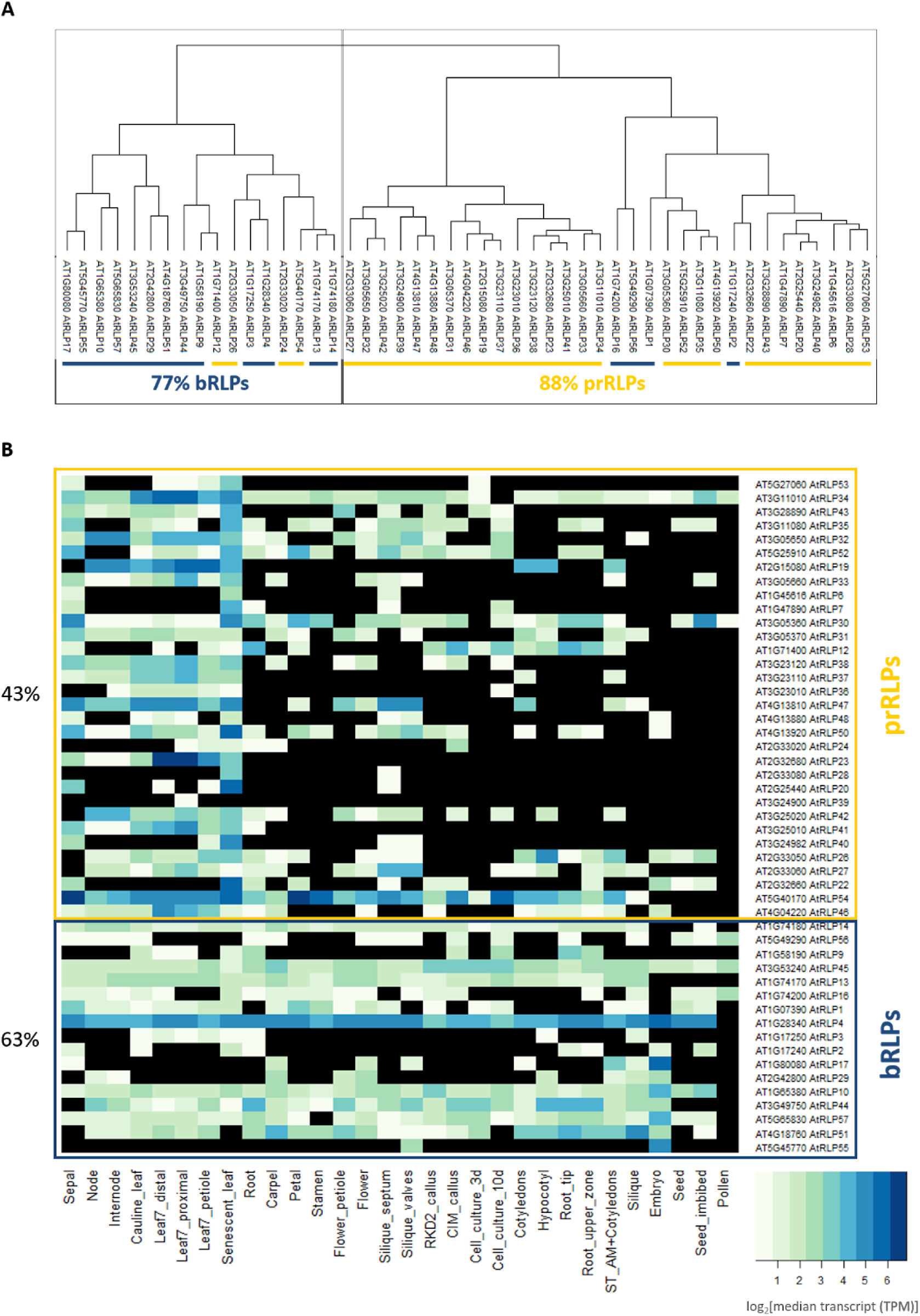
Transcriptomic clustering. (A) The dendrograms represent the hierarchical clustering of the transcripts of RLP-genes in various tissues after imputation of missing values (Mergner *et al.* 2020). (B) The heatmap shows the absence/presence of RLP transcripts in various uninduced tissues in Arabidopsis (Mergner *et al*. 2020) coloring indicates absence of transcript. Transcript abundance is indicated by the coloring code as log_2_ of median transcript (TPM, transcript per million). Black means no transcript was found. The presence of each gene transcript over all tested tissues was calculated and the average for each set (prRLPs and bRLPs) is shown in percentage. Boxed in yellow are prRLPs and in blue bRLPs.

For many RLPs expression data was only available for very few of the analyzed tissues. In Figure 3B, we therefore show a heatmap depicting the gene expression levels. Closer inspection shows that the fraction of RLPs with a detectable transcript differs significantly between the prRLPs and bRLPs, with a lower fraction detected in the prRLPs, 43% vs 63% (Student’s t-test, p=0.036). Thus, basal gene expression levels between RLP classes differ.

### Basal and pathogen-responsive RLPs cluster in the Proteome

Knowing that there are significant tissue specific differences in expression between the two RLP classes, we further investigated whether these differences can be observed at proteome level as well. When calculating clustering on protein abundance, we also observed a clustering of the pr- and bRLPs (Figure 4A).

**Figure 4:**
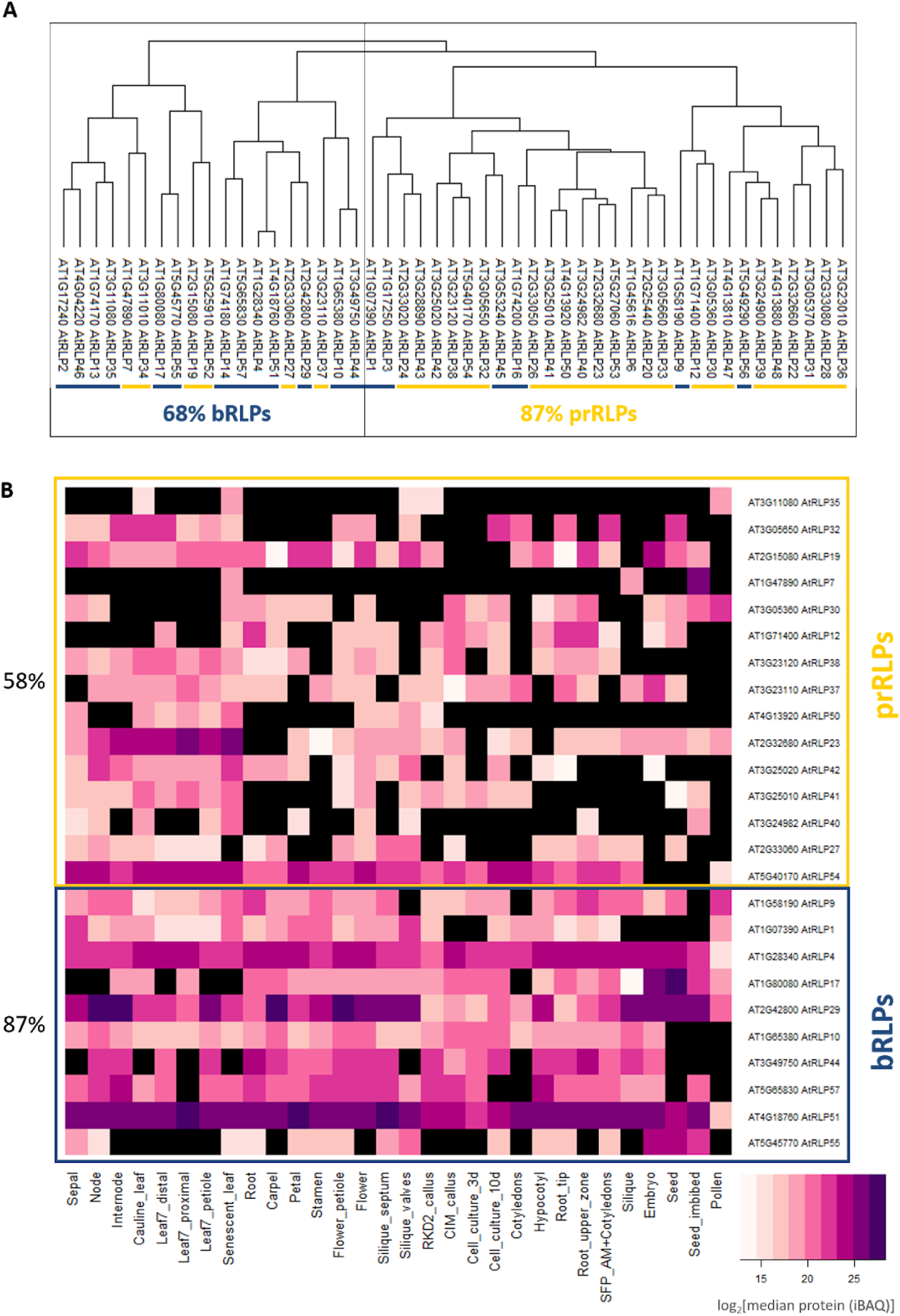
Proteomic clustering. (A) The dendrograms represent the hierarchical clustering of the translated RLP-proteins in various tissues after imputation of missing values (Mergner *et al.* 2020). (B) The heatmap shows the absence/presence of RLP protein in various uninduced tissues in Arabidopsis (Mergner *et al.* 2020). Protein abundance is indicated by coloring code as log_2_ of median protein (iBAQ). Black means no protein was found. The presence of each protein over all tested tissues was calculated and the average for each set (prRLPs and bRLPs) is shown in percentage. Boxed in yellow are prRLPs and in blue bRLPs.

Similar to the transcriptome data, the proteome data show significant differences between the fraction of RLPs present in the pathogen-responsive fraction (prRLPs 58%) versus the basal fraction (bRLPs 87%) (Student’s t-test, p=0,0005) (Figure 4B).

### Pathogen-responsive RLPs lie more often in clusters

It has been hypothesized that the physical location of defence associated genes, like those in the NLRs and RLP families, allows for more rapid evolution and recombination and that as such, these gene families evolved in clusters on the genome. Indeed genes in both families are often co-occurring and clustered on the genome (Andolfo *et al*. 2013), yet singleton RLPs have also been reported. In order to test whether prRLPs are more often occurring in clusters and other RLPs more often as singletons, we reassessed available genome annotation data and defined RLP clusters (table 1 and Figure 5). This shows that the fraction of clustered RLPs is higher in the pathogen-responsive RLPs (27 clustered, 7 singletons) than in the basal (7 clustered, 13 singletons) (*χ*^2^ test, p = 0.003).

**Figure 5:**
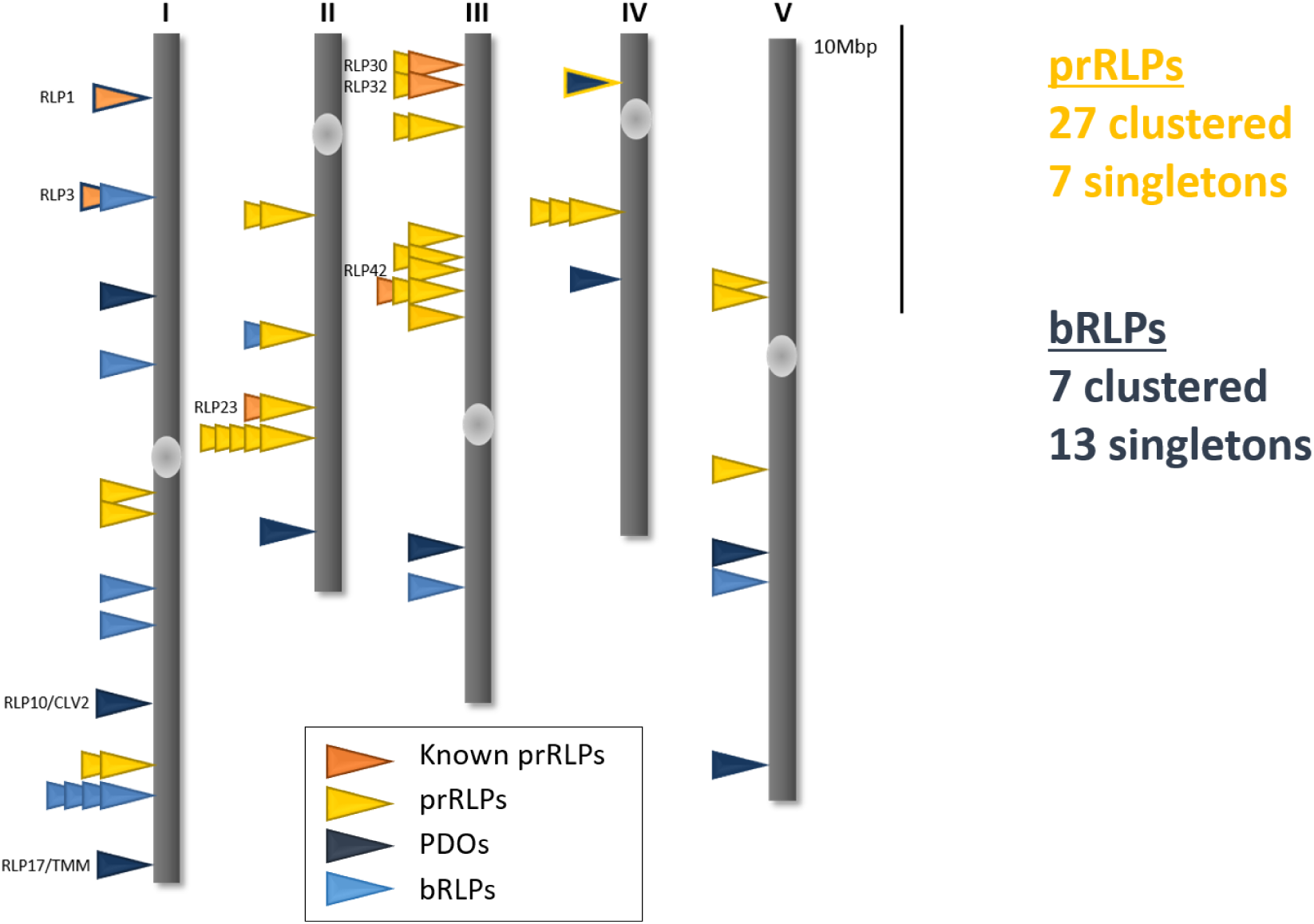
Genomic localization of RLPs. The genomic distribution is schematically depicted. 27 prRLPs are clustered and 7 are encoded as single genes, whereas 7 bRLPs are encoded in clusters and 13 as singletons (*χ*^2^-test, p=0.003). Known prRLPs are marked in orange, prRLPs in yellow, PDOs in dark blue and bRLPs in light blue. Figure is adapted from Tör *et al.* 2004.

**Figure S2:**
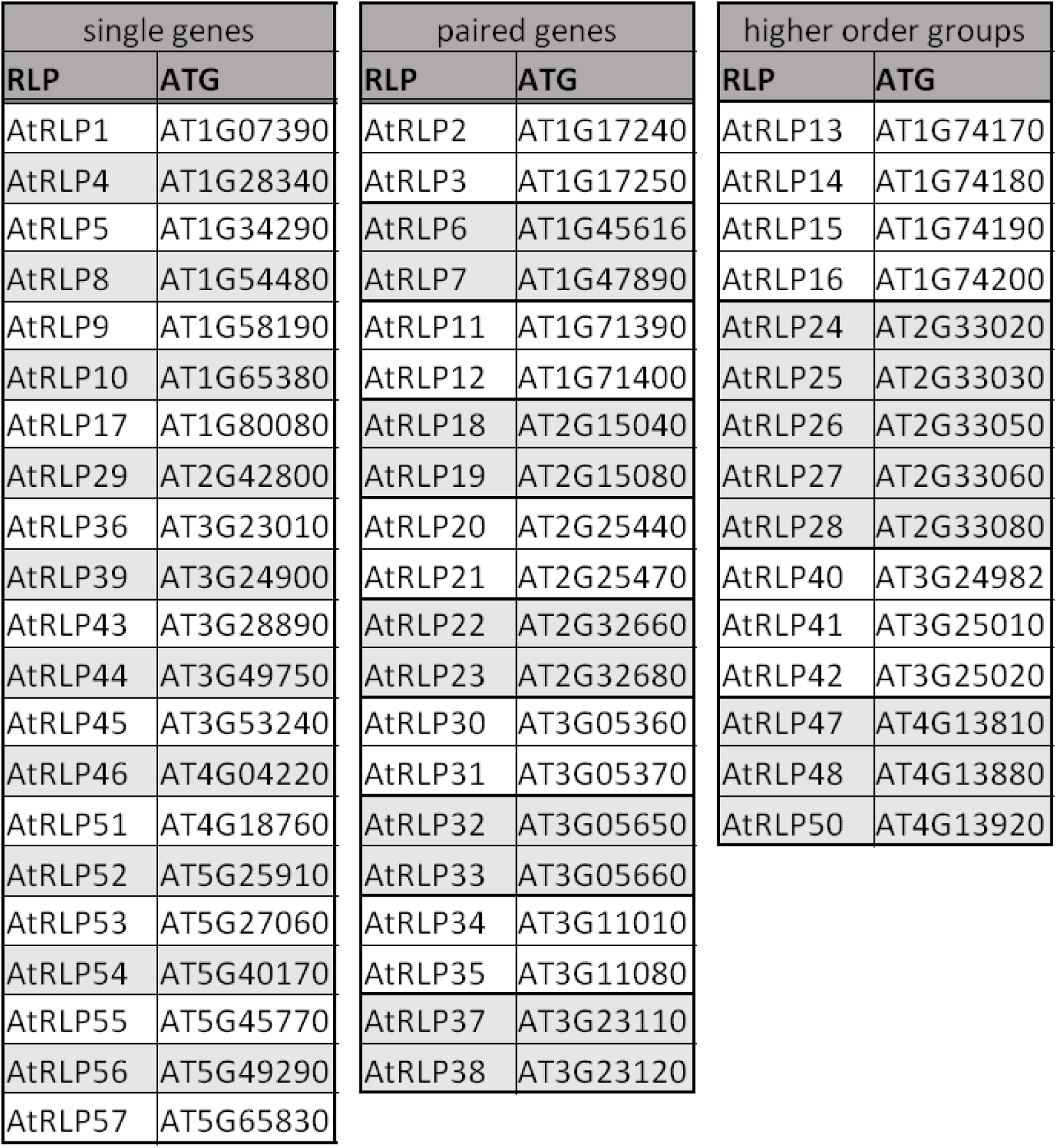
Genomic organization of RLPs in Arabidopsis.

### CNVs affect both classes of RLPs similarly

Gene clustering is expected to allow easier generation of copy number variation on affected chromosomal loci. In order to test whether defence RLPs differ significantly in CNV compared to other RLPs, we downloaded the CNV database generated by Zmienko *et al*. 2020, who defined CNVs as full as well as partial duplication of a gene or gene fragment. Interestingly, whereas CNVs are particularly widespread for NLRs (Van de Weyer *et al*. 2019), just over half of the RLPS (32) showed one or more CNV events. RLPs that occur in clusters are significantly more often affected by CNV events with 71% of clustered RLPs showing CNVs against only 40% of singleton RLPs (*χ*^2^-test, p = 0.05), thus clustering indeed seems to affect the potential for CNV in these genes. There is a tendency that indicates that prRLPs more often show CNV events (*χ*^2^-test, p = 0.14).

### RLPs show a broad range of single nucleotide polymorphisms (SNPs)

Now, knowing that pathogen-responsive RLPs are often found in clusters, and that this clustering might lead to an observed increase in CNV, we wanted to test whether pathogen-responsive RLPs are in general showing higher numbers of polymorphisms and higher signatures for positive or balancing selection. Analyzing the sequencing data from 1135 Arabidopsis accessions revealed that 22 out of 57 RLPs have no coding SNPs. 41% (14/34) of the prRLPs and 33% (7/21) of the bRLPS have no SNPs. Thus, the fractions are not significantly different (*χ*^2^-test, p = 0.87). Looking specifically in the clustered and non clustered RLPs revealed that no difference in the absence or presence of SNPs between RLPs which are encoded as a single gene or those in pairs or larger clusters (40% [8/20] vs 39% [14/34]).

Seeing that there are no significant differences between the total number of segregating sites, we looked whether other parameters are different between the two RLP classes and split the analyses in synonymous (e.g. not causing an amino acid change) and non synonymous (causing an amino acid change) SNPs. To our surprise, the total number of segregating sites and the nucleotide diversity measured as π/site are significantly larger in the bRLPs (Figure 6). Tajima’s D, which can be seen as a proxy for evolutionary pressure on the genes, is generally low in all RLPs. This is an indication of an abundance of rare alleles (singleton SNPs) being present and thus suggests purifying selection or a recent bottleneck. Low Tajima’s D values have been reported for the majority of genes in *A. thaliana* (Alonso-Blanco *et al.* 2016). Interestingly, whereas higher Tajima’s D would be expected for defence-associated genes under diversifying or balancing selection, there are only two RLPs with Tajima’s D values above 0, both of them belong to the bRLPs (RLP2, 0.822; RLP15 0.05). Lastly, we found no significant differences in the ratio of non-synonymous over synonymous polymorphisms between the two classes (Figure S3). Thus, it appears that on the level of DNA polymorphisms, prRLPs and bRLPs cannot be differentiated.

**Figure 6:**
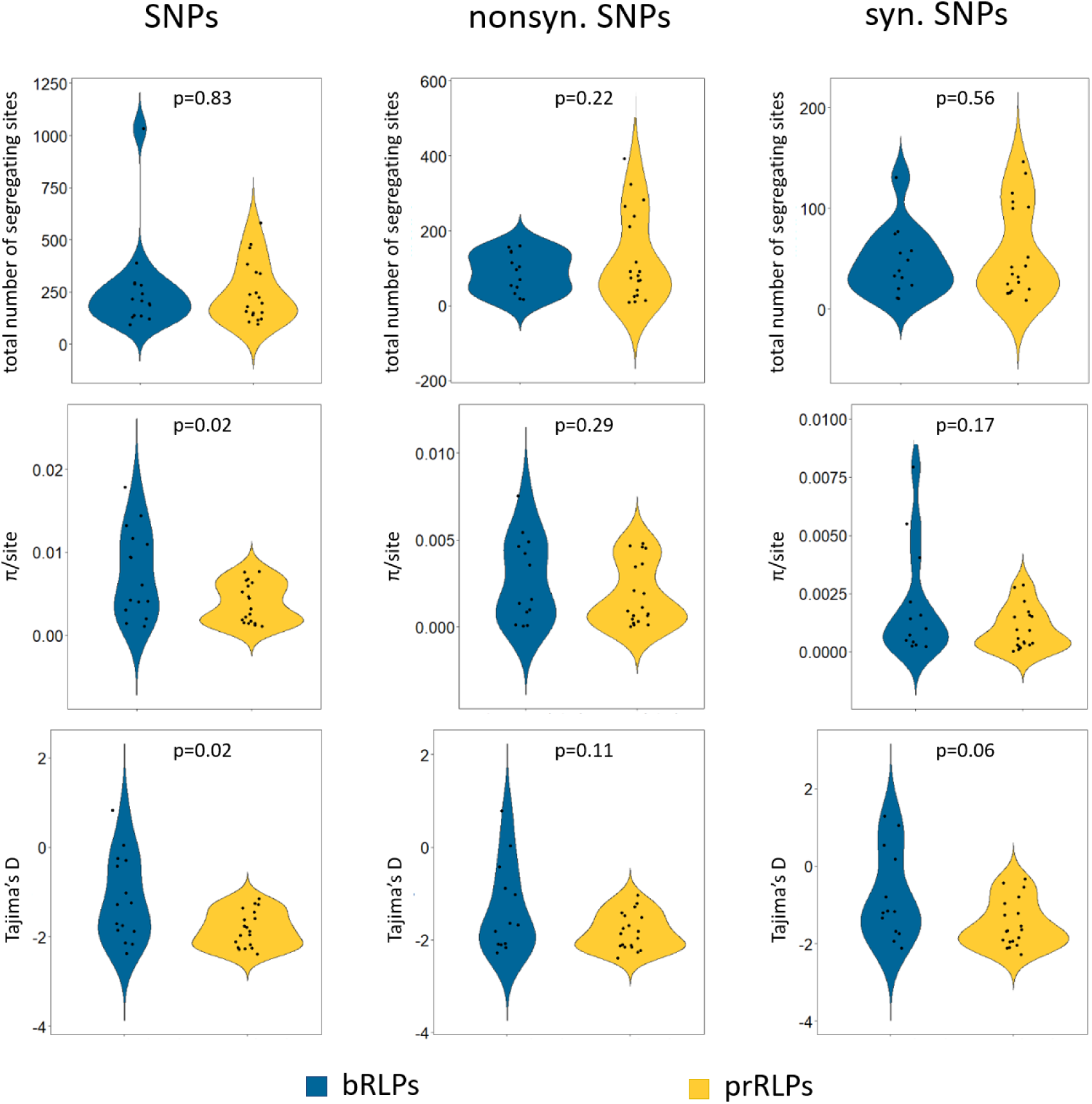
Single nucleotide polymorphisms (SNPs). We used the available sequence information of more than 1000 Arabidopsis thaliana accessions from the 1001 Genome project (Alonso-Blanco *et al.* 2016) and calculated the total number of segregating sites, π/site and Tajima’s D value for all SNPs, as well as nonsynonymous and synonymous SNPs using the *Popgenome package* (*Pfeifer et al.* 2014). P-values are calculated using Student’s t-test.

**Figure S3:**
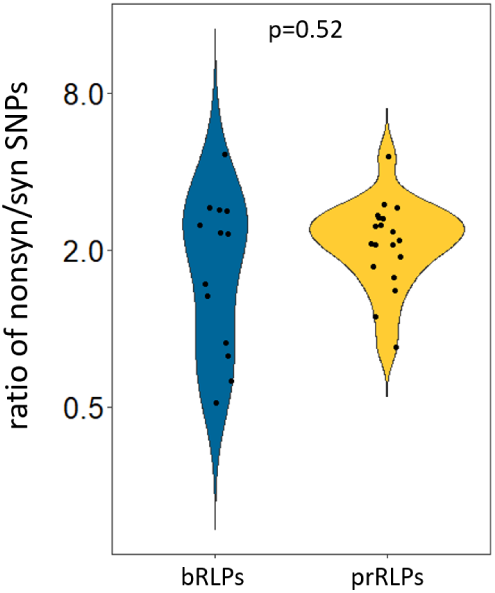
Ratio of nonsynonymous over synonyms SNPs on a logarithmic scale (y axis) as calculated with PopGenome for the bRLPs and prRLPs (x axis).

## Discussion

LRR-RLPs form a diverse gene family that has been associated with both developmental and defence associated processes. In this paper, we have combined publically available datasets to classify LRR-RLPs and predict putative roles in either defence or more basal, most likely development-associated processes.

The phylogenetic tree based on the C-terminal conserved domains C3 to F constructed by Wang *et al*. 2008a together with gene expression data collected by the Genevestigator database showed a very clear separation into LRR-RLPs upregulated (prRLPs) and not regulated (bRLPs) after various pathogen treatments.

This separation is further confirmed by analysis done by Fritz-Laylin *et al*. (2005). In this analysis the LRR-RLP sequences of *Arabidopsis thaliana* and rice were compared and based on different criteria, like homology and genomic localization a set of nine putative developmental orthologues (PDOs) was defined. This set includes the well-studied CLV2/RLP10 and TMM/RLP17 proteins, as well as RLP44 integrating cell wall surveillance with hormone signaling to control cell wall integrity and growth and to control cell fate in the root vasculature (Wolf *et al*. 2014; Holzwart *et al*. 2018). These 9 PDOs are all, except RLP46, not upregulated after pathogen treatment and cluster within the bRLPs.

Interestingly, our defined bRLPs are more closely related to tomato RLPs than to the Arabidopsis prRLPs (Kang and Yeom 2018), which form a unique clade, indicating that each species might have a unique set of receptors to fight off invading pathogens, but share commonalities in their basal processes.

RLPs lack an intracellular signalling domain and therefore need interaction partners for downstream signalling. The confirmed defence associated RLPs constitutively interact with the adaptor-kinase SOBIR1 and recruit BAK1 in a ligand-dependent manner (Albert *et al*. 2015, 2019). The signalling of those RLPs is SOBIR1-dependent, whereas the known PDOs (CLV2/RLP10, TMM/RLP17, RLP44) function independent of SOBIR1, but can be pulled down in overexpression experiments together with SOBIR1 (Liebrand *et al*. 2013). Two protein-interacting motifs are required for RLP-SOBIR1 interaction, which is a negatively charged stretched amino-acids in the extracellular juxtamembrane region and a GxxxG-motif in the transmembrane domain (Albert *et al.* 2019). Alignment of these regions showed that in most of the prRLPs both of the motifs are present, whereas in the bRLPs they are less common or completely absent.

We analysed the transcriptomic and proteomic expression profiles of the RLPs in 30 different samples, representing different tissues and different development stages (Mergner *et al*. 2020) and compared the prRLPs with the bRLPs. Both, the transcriptome and the proteome showed a separation of the bRLPs and prRLPs and it revealed furthermore that the bRLPs are more ubiquitous expressed and transcribed compared to the prRLPs.

Over the past years a number of studies have been published that aimed to identify receptors involved in early pathogen defence responses, so called pattern recognition receptors (PRRs). Our data show that all known RLPs that function as such PRRs, show similar patterns in the transcriptome and proteome data, especially the presence of the respective protein in an uninduced state. Constitutive presence of a cell surface receptor hence appears as a hallmark of PRRs and is a prerequisite to measure early responses to potential immunogenic elicitors from pathogens. Based on these combined data we can thus predict further RLPs that may act as PRRs and can potentially be identified using early immune response assays, as for example done by Zhang *et al.* 2013 or Albert *et al.* 2015. Expression data show upregulation of certain RLP-genes upon pathogen stimulus, indicating that those respective pathogens might harbour the immunogenic motif recognized by the respective PRR. This is true for already identified PRRs like RLP23, which recognizes nlp20, a 20 amino acid long peptide present in NECROSIS AND ETHYLENE PRODUCING (NEP)-LIKE PROTEINS (NLPs) (Böhm *et al.* 2014b). NLPs are widespread among bacteria, fungi and oomycetes and the expression data on Genevestigator shows that RLP23 is highly upregulated after treatment with pathogens harboring an NLP.

Our predictions can be found in table S1. All of those RLPs belong to the prRLPs, the protein is present in an uninduced state, and all, except RLP54 are encoded in a cluster of at least two genes, all typical attributes that we have shown to be typical for PRRs.

**Table S1:**
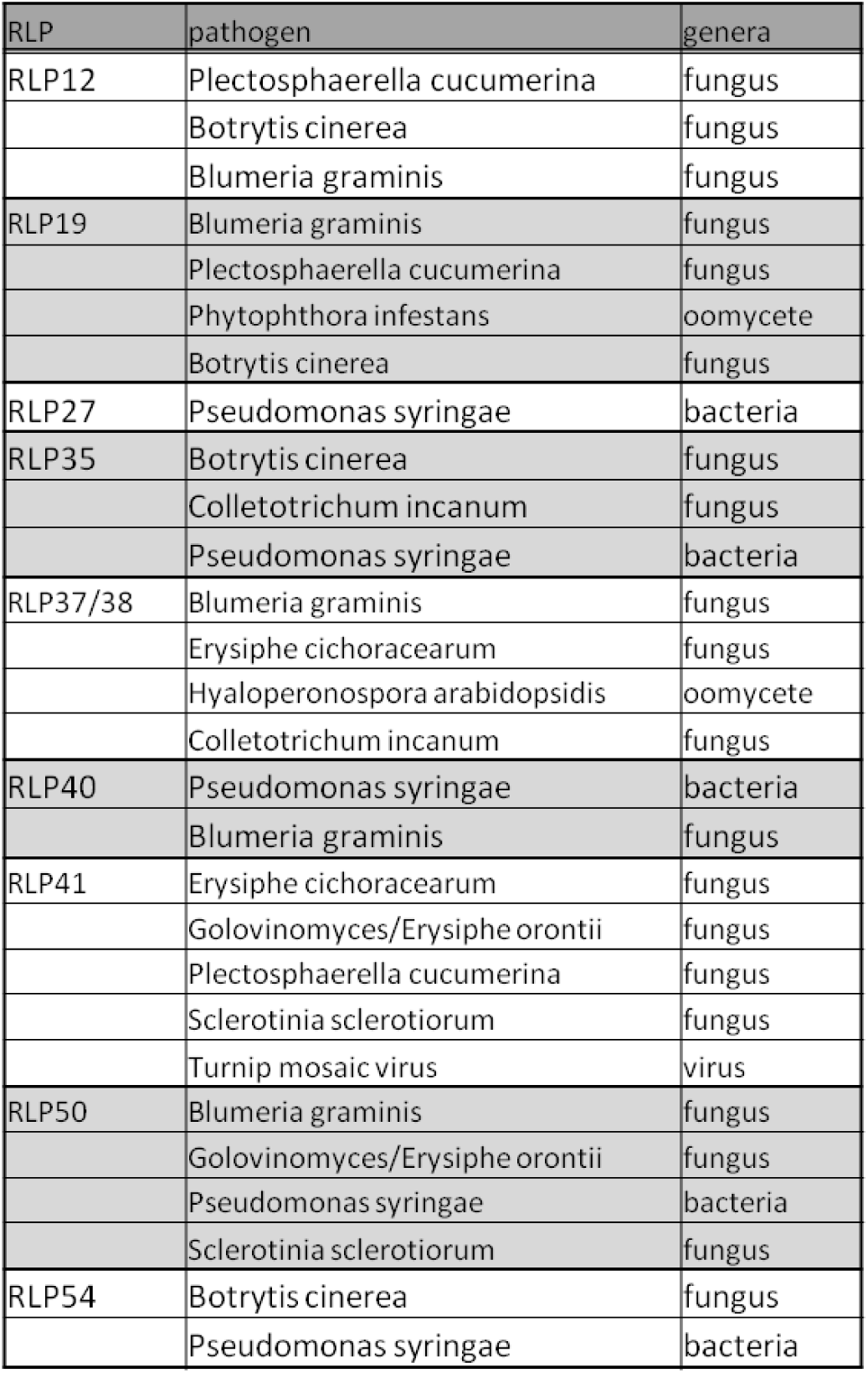
Putative PRRs with pathogens that might harbour an immunogenic motif, which is recognized by the respective RLP.

The gene expression atlas on Genevestigator (Hruz *et al.* 2008) further revealed that some RLPs (RLP4, 19, 21, 26/27, 32, 49/50, 53/34, 54) are likely to be targeted by bacterial effectors, as they showed a downregulation of gene expression after treatment with wild type bacterial strain and a strong upregulation after infection with bacterial strains deficient in effector secretion. Additionally, no RLP was upregulated after wounding, maybe indicating that RLPs are not able to sense damage-associated molecular patterns (DAMPs).

Besides the previously mentioned observation that the prRLPs seem to have evolved species specifically in *A. thaliana*, the bRLPs and prRLPs show a number of interesting genomic differences that illustrate possible differences in their evolutionary trajectory. We do not observe clear differences between the classes in terms of nucleotide diversity. This might be, because some of the bRLPs have dual roles (like CLV2) (Pan *et al.* 2016), or because the cluster of bRLPs also contains some defence associated RLPs. Nucleotide diversity and Tajima’S D differ between bRLPs and prRLPs, but this difference seems to be mainly driven by two highly diverse bRLP.

It might not come as a surprise that RLPs are generally not showing great diversity, as illustrated by the lack of SNPs in some and general low Tajima’s D, because developmental processes are assumed to be conserved and VDRs in *A. thaliana* are detecting conserved PAMPs. Yet, the stark differences in the amount of polymorphisms in some prRLPs as well as some bRLPs, might indicate specific roles for these more diverse RLPs.

We do find that prRLPs are significantly more often encoded in gene clusters than bRLPs and that prRLPs are more often affected by CVNs. Recently, such intragenic recombinations have been shown to play a major role in the maintenance of stable polymorphisms in an important NLR resistance gene against the pathogen *P. infestans* in the wild potato species *Solanum americanum* (Witek *et al*. 2020), as well as in the RLP locus Hcr9, conferring resistance against the fungal pathogen *Cladosporium fulvum* in wild tomato (Kahlon *et al*. 2020).

Overall, by combining several resources, we enhance the current knowledge of the RLP gene family in Arabidopsis. We firmly differentiate between a pathogen responsive sub family and a basal sub family, each with their own characteristics. This distinguishment will help researchers working on the biology of RLP and might form an interesting starting point for comparative studies in other plant species.

## Material and Methods

### Phylogenetic analyses

The phylogenetic tree of the Arabidopsis’ RLPs was taken from Wang *et al.* 2008a. The b- and prRLPs were assigned based on the gene expression data available on Genevestigator (Hruz *et al*. 2008). We checked for genes which were at least 2.5x upregulated with a p-value of 0.001 after infection with various pathogens. The used datasets were AT_AFFY_ATH1-0 and AT_mRNAseq_ARABI_GL-3. The phylogenetic tree of the RLPs from tomato and Arabidopsis was done by Kang and Yeom 2018.

### Domain alignment

The full-length protein sequence of all RLPs was aligned using muscle and the sequences were afterwards ordered manually to fit the phylogenetic tree and trimmed to the last LRR-domain, the extracellular juxtamembrane region, the transmembrane domain and the intracellular juxtamembrane region.

### Transcriptome and proteome data clustering

Both transcriptome and proteome data for 30 *Arabidopsis thaliana* tissues were obtained from the Arabidopsis Proteome project (Mergner *et al*. 2020). In this database, the transcriptome data were log normalised and transformed into fold changes over the median. Proteome data were transformed similarly. Missing data were imputed around the mean. For each of the two data sets the Pearson correlations were calculated between the values in R, using cor, followed by clustering using hclust (method = complete for proteome and ward.d2 for transcriptome) and plotting as.dendogram.

### Genomic clustering and CNV analysis

The genomic clustering is based on the analysis done by Tör *et al.* 2004 and the start sites of each gene. To test whether the observed number of bRLPs and prRLPs in clusters differed from expected values, we used the *χ*^2^-test (chisq.test) as implemented in the R stats package.

For CNV analyses, we used the CNV definition and the dataset as described in Zmienko *et al. 2020*. The genomic coordinates of the CNVs were extracted from the supplementary data and converted to bed format. Next we used bedtools intersect -wo to find overlapping regions. CNVs were counted for bRLPs and prRLPs and their distribution was compared with expected ratios using the *χ*^2^-test in R as described before.

### Genetic diversity analyses of 1001 genome data

The sequencing data for 1135 Arabidopsis accessions was downloaded from the 1001 genomes project homepage (https://1001genomes.org/) and the Col-0 reference genome was downloaded from (https://plants.ensembl.org/).

The nucleotide diversity statistics were calculated with the R package PopGenome (Pfeifer *et al*. 2014) using the functions diversity.stats and neutrality.stats. All statistics were calculated for all sites and separately for the synonymous and nonsynonymous sites using the subsites function. To obtain comparable π/site values, the π-value was divided by the gene length.

## Acknowledgements

We would like to thank all the authors of the used resources for publishing open access and sharing their data with the community, without it this whole work wouldn’t have been possible. Further, we would like to thank Prof. Dr. Ralph Hückelhoven for discussion and reading the manuscript.

## Notes

### Competing Interest Statement

The authors have declared no competing interest.

